# Genome-wide within-host evolution during prolonged SARS-CoV-2 infection in a person with advanced HIV infection

**DOI:** 10.64898/2026.10.07.756802

**Authors:** Karina Perlaza, Ishaan Dureja, Diane Havlir, Vivek Jain, Jeffrey Whitman, Monica Dayao, Bryan Castillo-Rojas, Julia Rosecrans, Melanie Ott, Carina Marquez, Joseph L. DeRisi, Taha Y. Taha

## Abstract

Within-host evolution during prolonged SARS-CoV-2 infection can be extensive in immunocompromised individuals, but the consequences of mutations that emerge and change in frequency over time remain difficult to predict. We characterized an XBB.1.5.10 infection lasting nearly 90 days in a person with advanced human immunodeficiency virus (HIV) infection. Genome-wide tracking of allele frequencies in samples collected on study days 43, 48, and 70 revealed a Spike N-terminal domain (NTD) deletion spanning residues 138-145 at frequencies of 33%, 80%, and 96%, respectively. A receptor-binding domain (RBD) deletion-insertion spanning residues 373-378 was detected on days 43 and 48 but not on day 70, while nsp12 C464Y was detected at approximately 8% on day 48 and 86% on day 70. We reconstructed representative Spike and non-Spike mutation profiles from the longitudinal samples and tested their effects on cell entry and viral RNA replication. The representative Spike construct for days 43 and 48 increased entry relative to XBB.1.5. Comparison with a matched construct lacking the receptor-binding domain deletion-insertion showed that the deletion-insertion enhanced entry, although it was not detected in the day 70 sample. The day 48 non-Spike profile supported the highest RNA replication among the tested replicons. Together, allele-frequency tracking revealed both transient mutations and mutations detected at progressively higher frequencies across the sampled time points. Functional testing showed that increased entry or RNA replication in vitro did not necessarily predict persistence in vivo.

**IMPORTANCE:** Severe immune suppression can allow SARS-CoV-2 to diversify within a single host, generating mutation combinations rarely observed during typical infections. In a person with advanced HIV infection, we tracked viral change across the genome and tested how representative mutation combinations affected cell entry and RNA replication. An eight-residue Spike deletion was detected at progressively higher frequencies across three sampled time points, whereas a separate Spike change enhanced entry in cell culture but was not detected in the final sequenced sample. In addition, mutations in the viral replication machinery, including substitutions previously reported in remdesivir-treated infections, changed in frequency over time. Our findings show that enhanced entry or replication measured in vitro does not necessarily predict whether the associated mutations will remain detectable later in an infection.

## INTRODUCTION

Impaired viral clearance in immunocompromised individuals can extend SARS-CoV-2 replication and create opportunities for extensive within-host diversification (1–6). Advanced HIV infection may be particularly permissive to prolonged viral replication and diversification. Ko et al. reported markedly greater within-host Spike diversity among people living with HIV and CD4 counts below 200 cells/µL than among people with higher CD4 counts or people without HIV infection (7). Longitudinal sampling can show how mutation frequencies differ across time points, but the functional consequences of these changes often remain uncertain.

Here, we characterized viral evolution during a nearly 90-day XBB.1.5.10 infection in a person with advanced HIV infection. We used longitudinal whole-genome sequencing to measure allele frequencies across three time points, including mutations below the consensus level, and examined linkage among nearby mutations in selected regions. We then tested representative Spike mutation combinations in cell-entry assays and representative non-Spike mutation profiles in RNA-replication assays.

## RESULTS

### Clinical course and treatment

We investigated SARS-CoV-2 evolution during a prolonged infection in a 49-year-old woman with advanced HIV infection (Fig. 1). SARS-CoV-2 was first detected approximately five weeks before hospitalization, and nirmatrelvir/ritonavir was prescribed but not taken. Study days denote days since first documented SARS-CoV-2 detection (day 0), as shown in Figure 1. The clinical course below uses hospital days, with hospital day 1 corresponding to study day 37. At hospital admission, she reported dry cough and non-bloody diarrhea and was febrile and tachycardic, with normal oxygen saturation on room air. SARS-CoV-2 testing remained positive (cycle threshold [Ct] 21), and chest computed tomography showed ground-glass opacities and small bilateral pleural effusions. She had restarted antiretroviral therapy (ART) approximately three weeks before admission. Her admission CD4 count was 14 cells/µL, and HIV RNA was approximately 500 copies/mL on hospital day 3, compared with approximately 500,000 copies/mL six months earlier.

**FIG 1.**
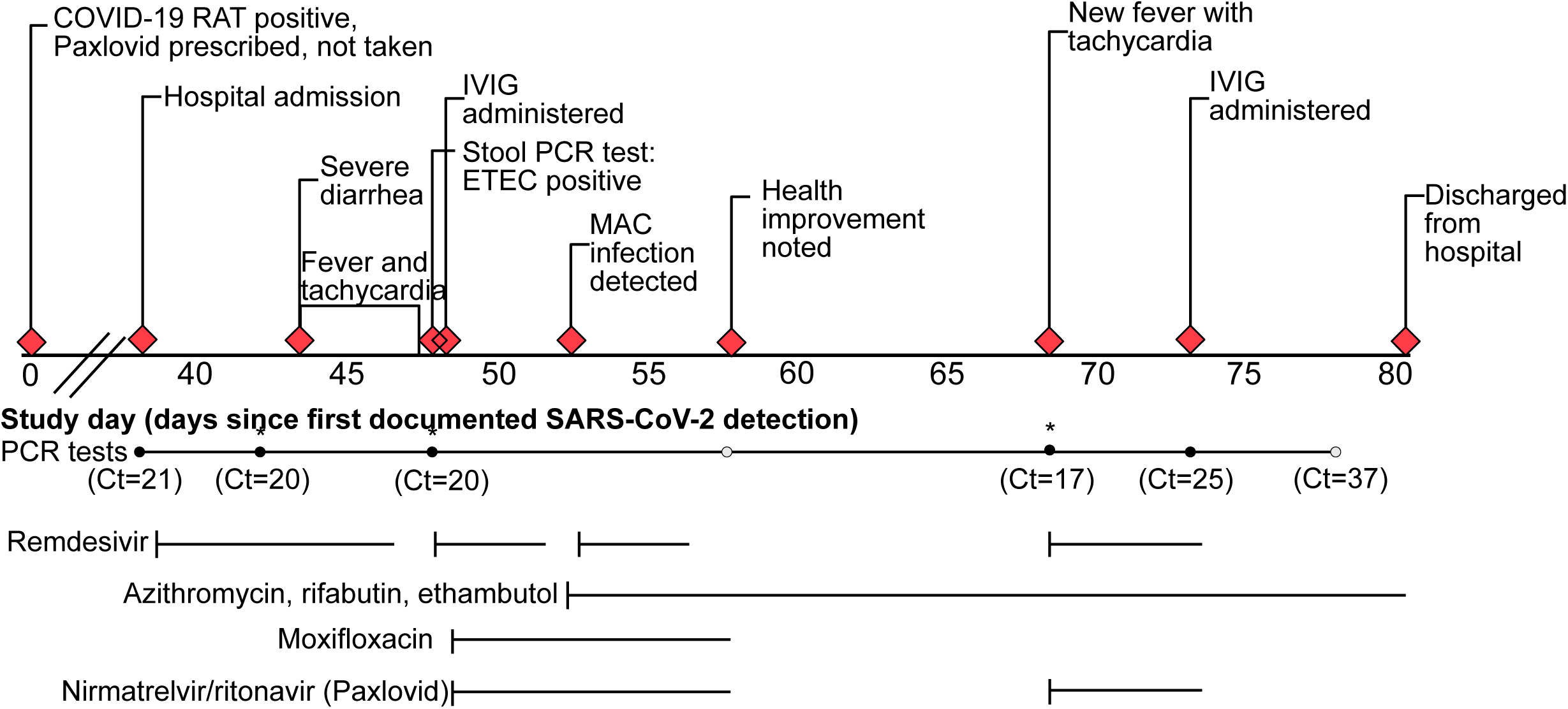
Clinical timeline of prolonged SARS-CoV-2 infection in a person with advanced HIV infection and disseminated MAC. Clinical events, antimicrobial and antiviral treatment courses, and SARS-CoV-2 testing results are shown as days since first documented SARS-CoV-2 detection. Day 0 denotes the first documented positive SARS-CoV-2 rapid antigen test. Hospital days in the text are numbered separately from admission, with hospital day 1 corresponding to study day 37. The nirmatrelvir/ritonavir course prescribed at first detection was not taken. Red diamonds indicate key clinical events. Filled circles indicate SARS-CoV-2 PCR-positive tests, and open circles indicate PCR-negative tests. Asterisks indicate nasopharyngeal swabs collected for viral genome sequencing on days 43, 48, and 70. HIV, human immunodeficiency virus, MAC, *Mycobacterium avium* complex, IVIG, intravenous immunoglobulin, PCR, polymerase chain reaction, Ct, cycle threshold.

During hospitalization, ART (darunavir/cobicistat/tenofovir alafenamide/emtricitabine plus dolutegravir) was continued, remdesivir was administered for persistent SARS-CoV-2 infection, and trimethoprim/sulfamethoxazole was given to prevent *Pneumocystis jirovecii* pneumonia. Persistent SARS-CoV-2 positivity prompted extended remdesivir treatment, with nirmatrelvir/ritonavir and intravenous immunoglobulin (IVIG) added on hospital day 11. Fever and diarrhea persisted, and stool PCR detected diarrheagenic *Escherichia coli*. Blood cultures were reported positive for *Mycobacterium avium* complex (MAC) on hospital day 16, prompting treatment with azithromycin, ethambutol, and rifabutin. SARS-CoV-2 PCR became negative on hospital day 20 after the initial antiviral course.

After initial improvement, fever and tachycardia recurred on hospital day 31 (study day 67), accompanied by renewed SARS-CoV-2 positivity (Ct 17). Remdesivir plus nirmatrelvir/ritonavir was restarted while persistent infection versus reinfection was considered. Antivirals were stopped on hospital day 36, when PCR remained positive (Ct 25), and a second IVIG infusion was administered. PCR remained weakly positive on hospital day 42 (Ct 37) and was negative by discharge on hospital day 44. HIV RNA was undetectable at discharge, and she continued ART and MAC treatment after returning home.

### Longitudinal viral sequencing and mutation frequencies

Nasopharyngeal swabs were collected 43, 48, and 70 days after first documented SARS-CoV-2 detection. Viral genome sequencing and phylogenetic placement confirmed that all three samples clustered within XBB.1.5.10, supporting within-host diversification rather than reinfection (Fig. 2A-B). The patient lineage was XBB.1.5.10, whereas XBB.1.5 served as the experimental reference for the functional constructs and assays described below. To summarize where coding variation accumulated, all amino acid changes detected across the three time points were aggregated, including both major and minor alleles passing the thresholding criteria (Fig. 2C). Most called variants were nonsynonymous and were distributed across multiple viral proteins, including Spike, other structural proteins, non-structural proteins, and accessory proteins (Fig. 2C-D).

**FIG 2.**
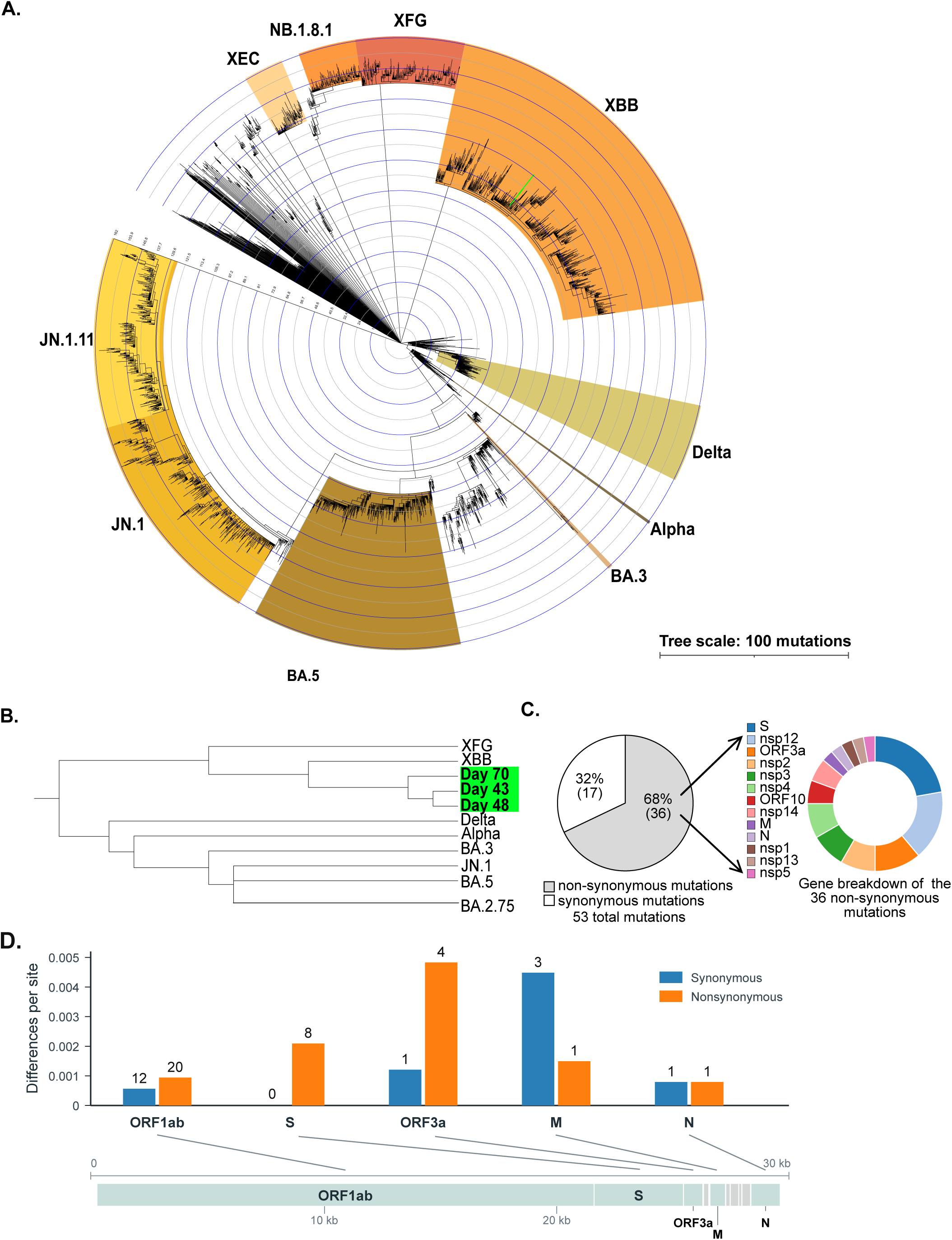
Phylogenetic placement and genome-wide mutation summary across longitudinal samples. (A) Reference maximum-likelihood tree of SARS-CoV-2 sequences colored by Nextclade clade assignment. The three patient sequences were placed onto the reference tree with Nextclade and cluster within clade 23A (XBB.1.5.10). (B) Expanded view showing that sequences from days 43, 48, and 70 form a distinct within-host clade. (C) Aggregate summary of variants detected across the three samples after quality filtering. The left chart shows synonymous and nonsynonymous variants, and the right chart shows the gene distribution of nonsynonymous variants. (D) Synonymous and nonsynonymous differences per site across the indicated viral genes. Numbers above the bars indicate the number of variants in each category.

We next tracked within-sample allele frequencies across days 43, 48, and 70 for nonsynonymous changes throughout the viral genome, including both major and minor alleles (Fig. 3). Variants were displayed if they reached an allele frequency of at least 10% at one or more sampled time points. To assess how frequently these same changes occurred outside this patient, we queried the 29 changes represented in Fig. 3 in public SARS-CoV-2 consensus sequences. During June 2023–June 2024, 23 of the 29 changes occurred in fewer than 0.1% of evaluable US sequences (Fig. S3, Table S2).

**FIG 3.**
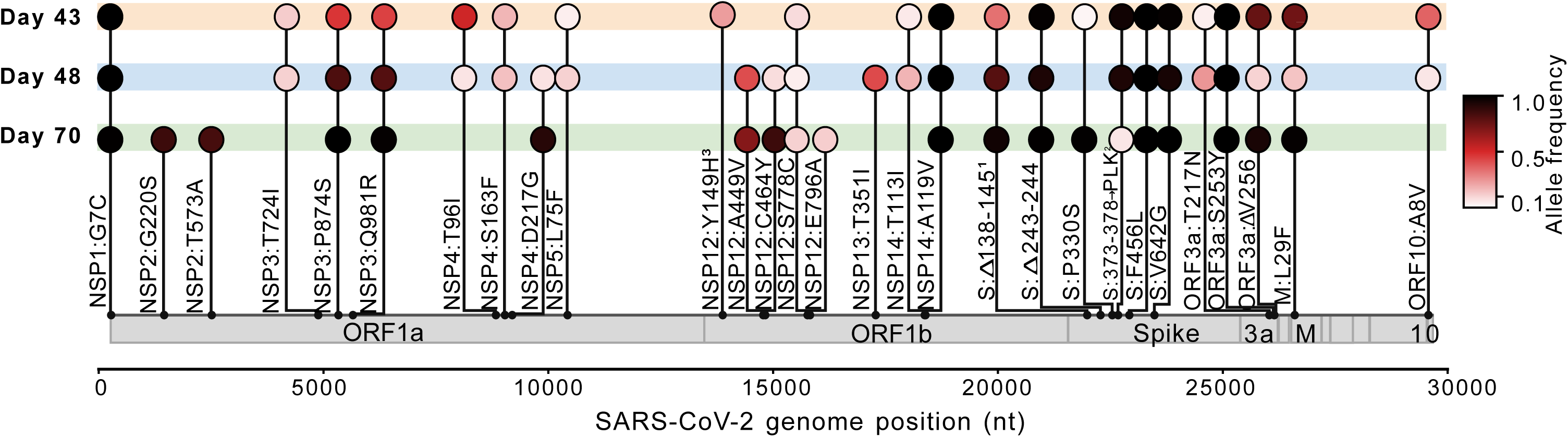
Genome-wide allele-frequency dynamics of within-host nonsynonymous variants across days 43, 48, and 70. Each circle represents a nonsynonymous variant at the indicated genome position and sampled time point. Circle color indicates allele frequency according to the scale at right. XBB.1.5 backbone variants were excluded. Both major and minor alleles are shown, and inclusion required an allele frequency of at least 10% at one or more sampled time points. Individual site-level calls were masked where locally linked sequence configurations were analyzed separately to avoid duplicate representation. Superscripts 1 and 2 identify the two locally phased Spike configurations analyzed in Fig. 4, and superscript 3 indicates insufficient coverage at day 70 for the marked site. Genome coordinates follow NC_045512.2, and gene boundaries are shown below the x-axis. Public-sequence frequencies of these changes are shown in Fig. S3 and Table S2.

### Spike variation in the NTD and RBD

Because several deletion and substitution calls occurred within the same short regions, we examined reads spanning these sites to determine which changes occurred together. This allowed us to distinguish the locally linked NTD and RBD sequence configurations shown in Fig. 4. The most prominent Spike changes occurred in the NTD region spanning residues 138-149 and the RBD region spanning residues 373-378 (Fig. 3). All three samples retained the XBB.1.5-defining L24S and Δ25-27 changes in the NTD. Additional changes in the patient sequences relative to XBB.1.5, detected at or near 100% in all three samples, included Δ243-244, F456L, and V642G (Figs. 3 and 4A).

**FIG 4.**
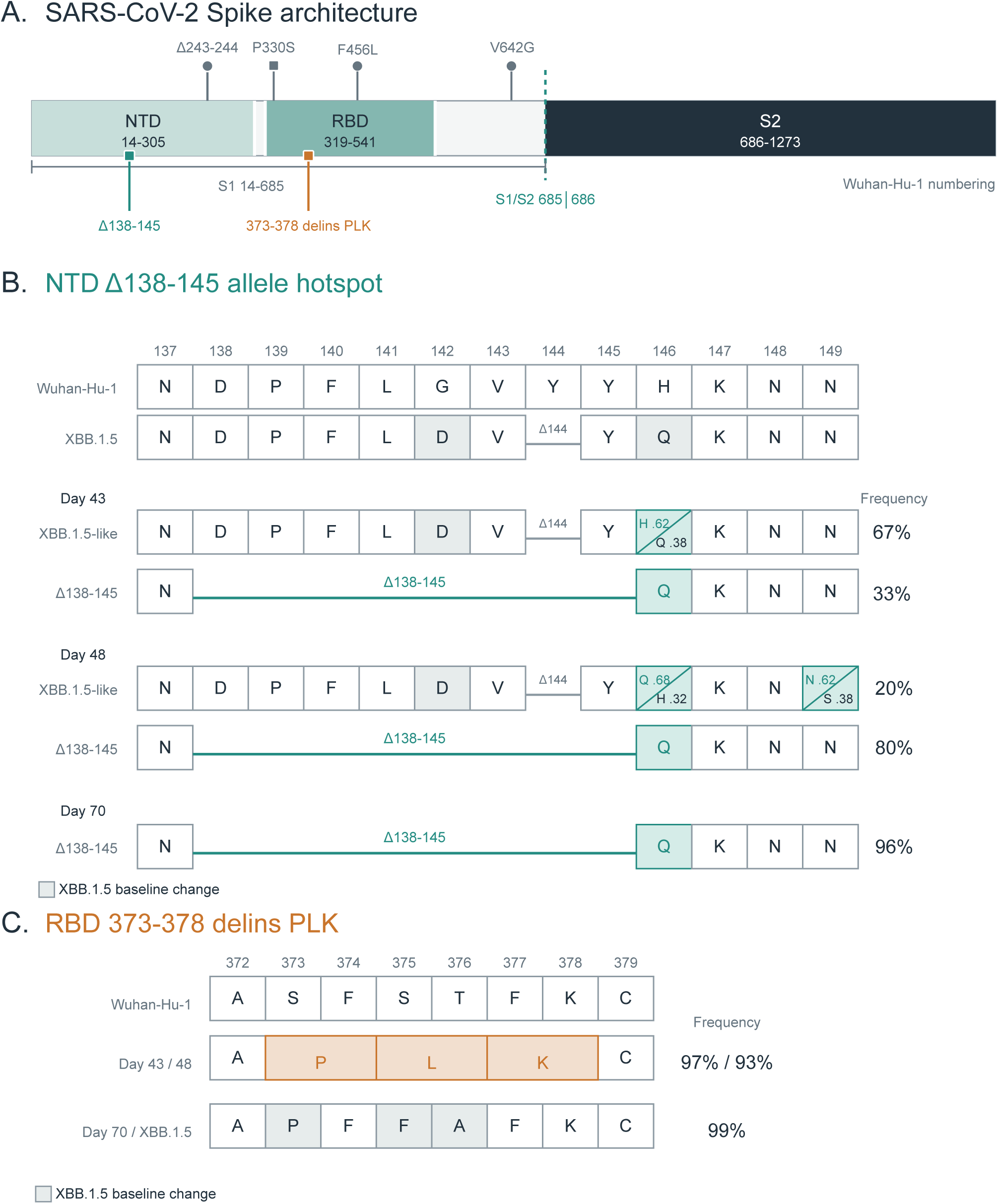
Locally phased analysis of NTD deletion and RBD deletion-insertion dynamics across longitudinal samples. (A) Spike domain map showing the positions of the principal patient-derived changes analyzed here. Residue numbering follows Wuhan-Hu-1. (B) Amino acid configurations spanning Spike residues 137-149 in the Wuhan-Hu-1 reference, XBB.1.5, and the locally linked NTD sequences detected on days 43, 48, and 70. The extended Δ138-145 configuration was detected at 33%, 80%, and 96%, respectively. Reads carrying this deletion consistently encoded Q146, whereas the XBB.1.5-like configuration showed mixed alleles at residues 146 and 149. (C) Amino acid configurations spanning Spike residues 371-379. The 373-378 delins PLK configuration was detected at 97% and 93% on days 43 and 48, respectively, but was not detected on day 70. The change is p.Ser373_Lys378delinsProLeuLys relative to Wuhan-Hu-1 and p.Phe374_Phe377delinsLeu relative to XBB.1.5. Shading indicates XBB.1.5 baseline changes as shown in the figure. NTD, N-terminal domain, RBD, receptor-binding domain.

The main NTD event was a deletion spanning residues 138-145. The deletion was detected at 33%, 80%, and 96% in samples collected on days 43, 48, and 70, respectively (Fig. 4B). Locally phased reads distinguished the extended Δ138-145 deletion configuration, which consistently encoded Q146, from the XBB.1.5-like configuration with mixed alleles at residues 146 and 149 (Fig. 4B). The 138-145 region overlaps recurrent deletion region 2 (RDR2, Δ141-146) (8) and the NTD antigenic supersite, a surface targeted by multiple neutralizing antibody classes (8–10).

A second prominent Spike change occurred in the RBD. The p.Phe374_Phe377delinsLeu change relative to XBB.1.5 (hereafter, the RBD delins) was present at 97% on day 43 and 93% on day 48 but was not detected in the day 70 sample (Fig. 4C). A separate RBD substitution, P330S, was the only RBD change unique to day 70 and was detected at 98.9%. The exact RBD delins product, APLKC between the retained A372 and C379 anchors, was not detected among 8,892,131 evaluable sequences in the worldwide, all-date search (Table S3A). Four US sequences collected in 2022 and assigned to the BA.1 lineage family contained the related local product APFKC, with phenylalanine at the position corresponding to leucine in the patient’s APLKC product (Table S3C).

Functional testing used single-round infectious SARS-CoV-2 replicon particles, preserving the SARS-CoV-2 structural-protein context and intracellular RNA replication rather than using a heterologous lentiviral or VSV backbone (37–38), as outlined in Fig. S4. For the entry assays, the day 43 and day 48 Spike mutation profiles were represented by a single construct because the same mutation set was selected at both time points (Fig. 5A). This construct differed from the persistent construct only by the RBD delins, allowing its effect to be tested on the patient-derived Spike background. Entry was measured in Vero cells stably expressing human angiotensin-converting enzyme 2 (ACE2) and human transmembrane serine protease 2 (TMPRSS2), referred to here as VAT cells, and in A549 cells stably expressing high levels of ACE2 (A549-ACE2 cells). In VAT cells, entry relative to XBB.1.5 increased from 1.57-fold with the persistent construct to 9.39-fold with the day 43/day 48 construct. In A549-ACE2 cells, entry increased from 1.13-fold to 10.98-fold (Fig. 5B). To test the RBD delins independently of the other patient-derived Spike changes, it was introduced directly into XBB.1.5 Spike and increased entry 8.24-fold in VAT cells and 52.14-fold in A549-ACE2 cells relative to XBB.1.5 (Fig. 5C). P330S was added to the persistent patient-derived changes to generate the day 70 Spike construct (Fig. 5A). Relative to XBB.1.5, the day 70 construct supported 1.00-fold entry in VAT cells and 1.18-fold entry in A549-ACE2 cells. These values were similar to those of the persistent construct (1.57-fold and 1.13-fold, respectively), indicating that P330S did not appreciably alter entry in either recipient cell type (Fig. 5B).

**FIG 5.**
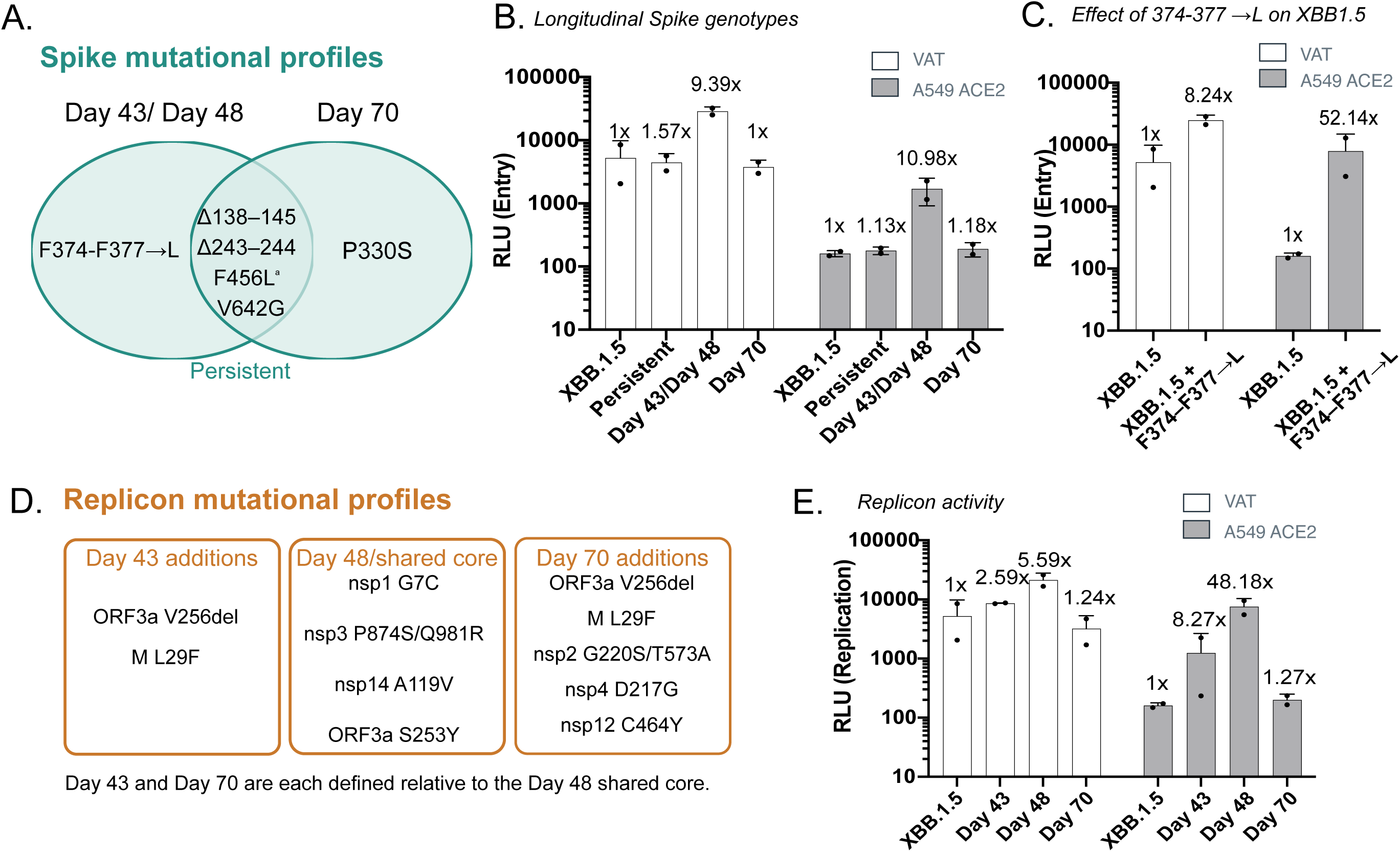
Representative patient-derived mutation profiles affect cell entry and support viral RNA replication. (A) Spike mutation profiles tested on the XBB.1.5 experimental backbone. The persistent profile contained Δ138-145, Δ243-244, F456L^a^, and V642G. The day 43/day 48 profile additionally contained the RBD delins, whereas the day 70 profile additionally contained P330S. ^a^F456L is an XBB.1.5.10-defining change absent from the XBB.1.5 experimental Spike backbone. (B) Entry of particles bearing the indicated Spike profiles into VAT and A549-ACE2 recipient cells. The selected day 43 and day 48 Spike profiles were identical and were tested as one construct. (C) Entry of particles bearing XBB.1.5 Spike with or without the RBD delins. (D) Patient-derived non-Spike mutation profiles incorporated into the day 43, day 48, and day 70 replicons. The day 43 and day 70 profiles are each defined relative to the day 48 shared core. (E) RNA replication supported by the indicated replicons in VAT and A549-ACE2 recipient cells. Values in panels B, C, and E were normalized within each replicate to the matched XBB.1.5 control. Bars show mean ± standard deviation of two replicate measurements per condition (n = 2). Complete construct compositions are provided in Table S1. VAT, Vero cells stably expressing human angiotensin-converting enzyme 2 (ACE2) and human transmembrane serine protease 2 (TMPRSS2), A549-ACE2, A549 cells stably expressing high levels of human ACE2. The assay design is shown in Fig. S4.

### Replication-complex allele frequencies and RNA replication

Replication-complex allele frequencies changed substantially between days 48 and 70. At day 48, nsp12 A449V and nsp13 T351I were present at nearly identical frequencies of 41.2% and 41.9%, raising the possibility that they occurred together on the same viral genomes. The sites were too far apart for individual sequencing reads to confirm this. By day 70, nsp12 A449V had risen to 69.0%, C464Y had increased from 8.1% to 85.7%, and C464F was no longer detected. Because coverage at nsp13 T351I was insufficient at day 70, we could not determine whether it persisted. nsp14 A119V was present at all three time points. The representative non-Spike profiles incorporated into each replicon are shown in Fig. 5D. Replicons containing the day 43, day 48, and day 70 profiles supported RNA replication at or above the XBB.1.5 reference level in both recipient cell types. The day 48 profile showed the highest RNA replication in both cell types, whereas the day 70 profile returned to near-reference levels (Fig. 5E).

### Additional variation occurs in ORF3a and the papain-like protease

ORF3a variation clustered within residues 253-263 of the cytoplasmic C terminus (Fig. 3). S253Y was observed at 100% allele frequency at all three time points, although coverage at this locus was limited, particularly at day 70. A three-nucleotide deletion encoding V256del fluctuated from 75.9% at day 43 to 10.2% at day 48 and 94.1% at day 70, also with lower coverage at the first and final time points. A larger overlapping 24-nucleotide deletion encoding Δ256-263 appeared at 8.9% on day 48.

Within nsp3, two substitutions localized to the papain-like protease (PLpro) domain: nsp3 P874S and Q981R, corresponding to PLpro P129S and Q236R, respectively (Fig. 3). Their allele frequencies rose nearly in parallel from 46.5% and 43.9% at day 43 to 81.8% and 80.2% at day 48, reaching 100% and 98.6% by day 70. Their nearly parallel frequencies prompted us to assess linkage, but the sites were too far apart to resolve on individual reads or read pairs. These variants were not functionally tested here, and their effects remain unknown.

## DISCUSSION

Across the three sampled time points, allele-frequency tracking distinguished mutations that persisted, increased, or were no longer detected in Spike and the replication-transcription complex. Functional testing showed that increased entry or replication in vitro did not necessarily predict whether the associated mutations would remain detectable at a later time point.

The Δ138-145 NTD deletion was detected at 33%, 80%, and 96% in the samples collected on days 43, 48, and 70. The deletion lies within recurrent deletion region 2 and the NTD antigenic supersite, a surface targeted by multiple neutralizing antibody classes and repeatedly altered across lineages (8–10). Although changes in this region are frequently associated with antibody escape, Ko et al. detected selected NTD mutations in individuals without detectable serum spike binding (7), suggesting that antibody pressure may not fully explain their emergence. NTD deletions may therefore affect viral properties other than antibody recognition. Similar but nonidentical deletions have been observed in population-level lineages. Yu et al. characterized a Brazilian isolate with an NTD deletion spanning residues 136-144 that retained fusion and entry despite the large deletion (11). Related large NTD deletions have appeared in the highly divergent BA.2.87.1 and BA.3.2 lineages (12–13).

V642G was present at or near 100% at each sampled time point. V642 lies near subdomain 2 (SD2), the 630 loop, and the fusion peptide proximal region (FPPR), which are implicated in conformational regulation and interprotomer packing (14–15). Although its functional effect remains uncharacterized, V642G has been observed alongside NTD deletions in at least three independent prolonged infections beyond the present study: with Δ144 in a lung transplant recipient with lower-airway disease on a BA.5 background (16), with Y144del and RBD deletions at residues 374-376 in an oncohematological patient with a BA.2 infection (17), and with large NTD deletions in a patient with advanced HIV infection and disseminated MAC infected with BA.2 (18). Thus, V642G has accompanied NTD deletions in four prolonged infections involving different viral lineages and causes of immunosuppression. BA.3.2 (“Cicada”) also shows extensive Spike divergence, including Δ136-147 and V642G (13) (Fig. S1). Whether V642G modifies the effects of these NTD deletions remains unknown.

The exact RBD delins was not identified in the public sequences surveyed, supporting its exceptional rarity within the available dataset (Table S3A), although similar deletions in this region have been reported. Rennick et al. described a probable B.1.1.7 transmission cluster carrying a nine-nucleotide deletion of residues 375-377 (19). When introduced into a Spike expression construct, Δ375-377 preserved syncytium formation but abolished binding of CR3022, an antibody recognizing a conserved cryptic RBD epitope (20). Ruotolo et al. also reported a high-frequency deletion spanning residues 374-376 during prolonged BA.2 infection in an immunocompromised patient (17). Here, the RBD delins increased entry both on the persistent patient-derived Spike background and when introduced directly into XBB.1.5 Spike, yet it was not detected in the day 70 sample. The only RBD change unique to day 70 was P330S, detected at 98.9%. Adding P330S to the persistent Spike background did not appreciably alter entry in either recipient cell type. The AlphaFold 3 model containing the RBD delins showed an apparent reorganization and reduction of local contacts between adjacent protomers (Fig. S2). Together, these observations show that increased entry in the tested cell systems did not predict whether the RBD delins remained detectable at a time point sampled later.

Between days 48 and 70, the predominant nsp12 alleles changed: A449V increased, C464Y rose from low frequency to dominance, and C464F was no longer detected. A449V and nsp13 T351I occurred at similar frequencies at day 48, but insufficient day 70 coverage prevented us from determining whether T351I persisted. C464Y and A449V have each been reported in remdesivir-treated immunocompromised infections (16, 21–23), although their co-occurrence has not been described to our knowledge. C464Y also emerged in multiple independent remdesivir-selected cultures (24). In the Igari et al. prolonged infection, low-frequency C464Y and C464F reached only 5% and 3% before fading (16), whereas C464Y became dominant here. In a separate prolonged infection, C464Y returned to fixation by the start of a second remdesivir course after a remdesivir-free interval (21), suggesting that continuous drug pressure was not required for its re-expansion. The persistent nsp14 A119V background may also be relevant because nsp14 exonuclease activity influences replication fidelity and viral fitness (25). These observations motivate testing whether nsp12 substitutions interact with contemporaneous nsp13 and nsp14 changes or alter remdesivir susceptibility.

ORF3a also varied within its cytoplasmic C terminus, although the functional effects of these changes were not tested (26–27). P129S and Q236R within PLpro are each individually tolerated for protease activity in deep mutational scanning (28), but their co-occurrence has not been reported to our knowledge. Interestingly, both residues are also present in the endemic human alphacoronavirus 229E (28). PLpro antagonizes innate immunity through deISGylation of host proteins (29–30), however, the functional consequence of this combination of mutations in the context of SARS-CoV-2 remains uncharacterized.

### Study limitations and future directions

Distinct viral genotypes have been identified across upper- and lower-respiratory-tract samples during prolonged infection (31), indicating that nasopharyngeal sampling may capture only part of the within-host viral population. In addition, our short-amplicon sequencing linked nearby changes but could not determine which distant mutations occurred together across complete viral genomes. Future studies combining near-complete viral-genome sequencing with host RNA profiling in individual infected nasal cells could resolve complete viral genotypes, associate them with specific cell types or states, and determine whether multiple genotypes occupy the same cell. Linked mutations may produce combined phenotypes not predicted from either mutation alone, while distinct viral genomes within a cell may complement or interfere with one another. The single-round infectious SARS-CoV-2 replicon system used here could then reconstruct candidate mutation combinations alone or in defined mixtures and test their effects on entry, replication, and cellular responses.

Longitudinal SARS-CoV-2 genomic characterization in this immunocompromised individual revealed substantial within-host variation across Spike, the replication-transcription complex, and host-interacting proteins. Rapid evolution and viral diversification in this setting may generate mutation combinations not yet observed in general circulation (5–6, 32). In this case, coupling genotypic characterization with functional assessment of entry and replication showed that enhanced entry or replication in vitro did not necessarily predict persistence within the host. Continued genomic and functional study of prolonged infections in immunocompromised patients may provide insight into SARS-CoV-2 evolutionary dynamics beyond what can be obtained from laboratory-based selections alone.

## MATERIALS AND METHODS

### Study design, clinical data, and sample collection

This study examined longitudinal SARS-CoV-2 evolution in one person with advanced HIV infection. Clinical events, laboratory results, and antiviral and antimicrobial treatments were abstracted from the medical record. Study days were defined as days since first documented SARS-CoV-2 detection, with the first positive test designated day 0. Hospital days were numbered separately, with admission designated hospital day 1 (study day 37). Remnant deidentified nasopharyngeal swabs collected on days 43 (TP1), 48 (TP2), and 70 (TP3) were used for viral genome sequencing under research protocols approved by the UCSF Human Research Protections Program.

### Viral RNA extraction and ARTIC amplicon sequencing

Total nucleic acid was extracted from nasopharyngeal swab material on an Agilent Bravo automated liquid-handling platform using a workflow adapted from the Quick-DNA/RNA Pathogen MagBead kit (Zymo Research). Complementary DNA was generated with LunaScript RT SuperMix. Two pools of overlapping approximately 275-nucleotide amplicons were generated with the UCSF CAT COVID-19 Tailed 275-bp v3 ARTIC protocol, based on the ARTIC Network v3 short-amplicon Illumina workflow (33). The two pools were combined, diluted, and indexed with 12-nucleotide TruSeq i5/i7 primers in a seven-cycle indexing PCR. Indexed libraries were pooled, purified with a 0.8× solid-phase reversible immobilization (SPRI) bead cleanup, assessed with an Agilent TapeStation D5000 assay, and sequenced on an Illumina platform using 2 × 147-bp paired-end reads. This tiled-amplicon approach follows a workflow previously used for SARS-CoV-2 genomic surveillance by the DeRisi laboratory (34).

### Read processing, variant calling, and annotation

Paired-end reads were processed with the CZ ID SARS-CoV-2 Consensus Genome Pipeline v3.4.18 (35). After adapter and quality trimming, reads were aligned to the Wuhan-Hu-1 reference genome (GenBank MN908947.3) with minimap2 v2.17-r941. Unmapped reads and alignments with mapping quality below 20 were removed with samtools. ARTIC primer intervals were then soft-clipped with iVar v1.3.1 (36) using the protocol-specific artic_v3_short_275_primers.bed coordinate file, a 2-nucleotide primer-position search offset, and retention of reads for which no primer was identified. The resulting sorted, primer-trimmed binary alignment map (BAM) files were used for downstream variant analysis.

Variants were called independently for each sample with FreeBayes in haploid pooled-continuous mode using a minimum base quality of 25, at least two alternate observations (AO), and a minimum alternate allele fraction of 0.002. Here and below, AO denotes alternate observations and DP denotes total read depth. Calls were normalized and decomposed into biallelic records with bcftools. Candidate single-nucleotide variants were tagged StrandDiscordant when AO was at least 10, the alternate allele fraction (AO/DP) was below 0.95, coverage was at least 50 observations in each orientation, and alternate support was no more than two observations in one orientation but at least 10 in the other. Tagged variants were excluded from the main callset. The main callset otherwise required AO of at least 10, total read depth of at least 30, and AO/DP of at least 0.02.

Variants were annotated with snpEff using the NC_045512.2 database. Per-base coverage was calculated with samtools, and uncalled positions with DP below 30 were classified as low coverage. Alleles were merged across time points into a coverage-aware frequency matrix and classified as called, low coverage, or not called. Open reading frame 1ab (ORF1ab) substitutions were mapped to the corresponding nonstructural proteins. Alleles detected at only one time point were retained at frequencies of at least 7.5%, whereas alleles detected at multiple time points were retained if they reached at least 5% at any time point. Predefined Spike deletion regions analyzed separately were excluded from the single-site matrix. XBB.1.5 backbone substitutions and manually resolved locally linked features were incorporated into the final analysis matrix.

### Manual review and local linkage of sequence changes

Selected calls, particularly complex variants and nearby linked changes, were manually reviewed in Geneious Prime 2023.1.1. The primer-trimmed BAM files generated by CZ ID were imported into Geneious and assembled against MN908947.3 for visualization. Read alignments were inspected to verify call quality, harmonize alternative representations of complex variants, and determine whether constituent changes co-occurred on individual reads or read pairs. This analysis established linkage only within the span of the amplicon reads or read pairs and did not establish linkage among distant mutations across complete viral genomes.

### Consensus sequences, lineage assignment, and phylogenetic placement

Consensus genomes were generated against Wuhan-Hu-1 (MN908947.3) with iVar as part of the CZ ID SARS-CoV-2 Consensus Genome Pipeline v3.4.18. Clade assignment and phylogenetic placement were performed with Nextclade. The reference tree was built with IQ-TREE2 under a maximum-likelihood model. Patient query sequences were placed onto the reference tree with Nextclade’s local greedy tree-building algorithm, and the tree was colored by Nextclade clade assignment.

### Selection of representative mutation profiles

Representative Spike and non-Spike mutation profiles were selected from the longitudinal allele-frequency data for functional testing. The constructs tested predominant mutation combinations selected at each time point and were not intended to reconstruct every viral genotype or the measured mixture of genotype frequencies in the samples. Because the amplicon data did not establish linkage among distant mutations, inclusion of changes in the same construct does not demonstrate that all of those changes occurred together on a single patient-derived genome.

For the entry assay, patient-derived Spike changes present at 30% or higher in all three samples were included in a persistent Spike background. Changes reaching at least 70% on day 43 or day 48 were added to this background. The selected day 43 and day 48 Spike mutation sets were identical and were therefore represented by one construct. P330S was added to the persistent background to generate the day 70 Spike construct. The receptor-binding-domain deletion-insertion was also introduced alone into XBB.1.5 Spike. For the RNA-replication assay, separate day 43, day 48, and day 70 non-Spike profiles were generated from the selected patient-derived changes at each time point. Complete XBB.1.5 backbones and the additional changes in every tested construct are listed in Table S1.

### Public sequence frequency analysis

Public SARS-CoV-2 consensus sequences were queried using open CoV-Spectrum/LAPIS (https://lapis.cov-spectrum.org/open/v2, October 1, 2026, data version 1790784010). US comparisons covered June 1, 2023–June 30, 2024 and September 1, 2025–September 30, 2026. Additional RBD searches included all available US dates and a worldwide exact-product search. Frequencies were calculated as matching sequences divided by sequences with unambiguous calls at the required positions, multiplied by 100. Query definitions and counts are provided in Tables S2–S3. September 2026 data were provisional.

### Spike expression plasmids and SARS-CoV-2 replicons

Spike expression plasmids and Spike-deficient SARS-CoV-2 replicons were assembled with pGLUE, a modular plasmid-based system for building SARS-CoV-2 genomes by Golden Gate assembly (37–38). The XBB.1.5 Spike expression plasmid (laboratory identifier SV4 XBB.1.5) and the XBB.1.5 Spike-deficient replicon (laboratory identifier pBACv6 XBB.1.5 s rep) used the same XBB.1.5 backbones described previously (38). In the replicon, the Spike coding region is replaced by secreted nanoluciferase and enhanced green fluorescent protein separated by a self-cleaving P2A peptide. The resulting single-round infectious particles contain the SARS-CoV-2 structural proteins Spike, membrane, envelope, and nucleocapsid, presenting Spike within a SARS-CoV-2 structural-protein context. The replicon retains intracellular viral RNA replication but cannot sustain subsequent rounds of infection without Spike complementation. Full backbone substitutions relative to Wuhan-Hu-1 and all added patient-derived changes are provided in Table S1.

### Replicon entry and RNA-replication assays

Replicon particle production and single-round infection assays were performed as described previously (38), with the assay design summarized in Fig. S4. Briefly, baby hamster kidney 21 (BHK-21) producer cells were transfected with the indicated replicon, Spike expression plasmid, and nucleocapsid expression plasmid. Supernatants were collected 72 h after transfection and filtered through 0.45-µm polyethersulfone AcroPrep Advance filter plates (Pall). Particle-containing supernatants were applied to Vero cells stably expressing human ACE2 and TMPRSS2 (VAT cells) or to A549 cells stably expressing high levels of human ACE2 (A549-ACE2 cells). Secreted nanoluciferase was measured 24 h after infection.

For entry comparisons, the pBACv6 XBB.1.5 Spike-deficient replicon was held constant while the indicated Spike expression plasmid was varied. For RNA-replication comparisons, SV4 XBB.1.5 Spike was held constant while the indicated replicon was varied. Values were divided within each replicate by the matched XBB.1.5 control without background subtraction. Data are shown as mean ± standard deviation of two replicate measurements per condition (n = 2). No inferential statistical tests were performed.

### Structural modeling

XBB.1.5 Spike and XBB.1.5 Spike containing the RBD delins (p.Ser373_Lys378delinsProLeuLys relative to WA1) were predicted as homotrimers using AlphaFold Server with default settings and five seeds (39). The top-ranked model (model_0) for each sequence was visualized in UCSF ChimeraX (40), and the region surrounding residues 373-378 near the interface between chains A and B was examined. Local atomic contacts were displayed as dashed pseudobonds using the ChimeraX Contacts tool and its default van der Waals-overlap criterion (overlap cutoff, −0.4 Å, atom pairs separated by four or fewer covalent bonds were excluded). Residue numbering follows the WA1 reference sequence.

### Data presentation

Allele frequency was calculated as AO/DP after application of the quality and coverage filters described above. Functional assay values were normalized within each replicate to the matched XBB.1.5 control and are presented as mean ± standard deviation of two replicate measurements per condition (n = 2).

## Competing Interests Statement

T.Y.T. and M.O. are inventors on a patent application filed by the Gladstone Institutes that covers the use of pGLUE to generate SARS-CoV-2 infectious clones and replicons. M.O. is a cofounder of DirectBio, Inc and on the SAB for Invisishield Technologies LTD. All other authors declare no competing interests.

## Funding

We gratefully acknowledge support from the Roddenberry Foundation, P. and E. Taft, Schmidt Futures, Gordon and Betty Moore Foundation, and the James B. Pendleton Charitable Trust to M.O..

## SUPPLEMENTAL FIGURE LEGENDS

**FIG S1.**
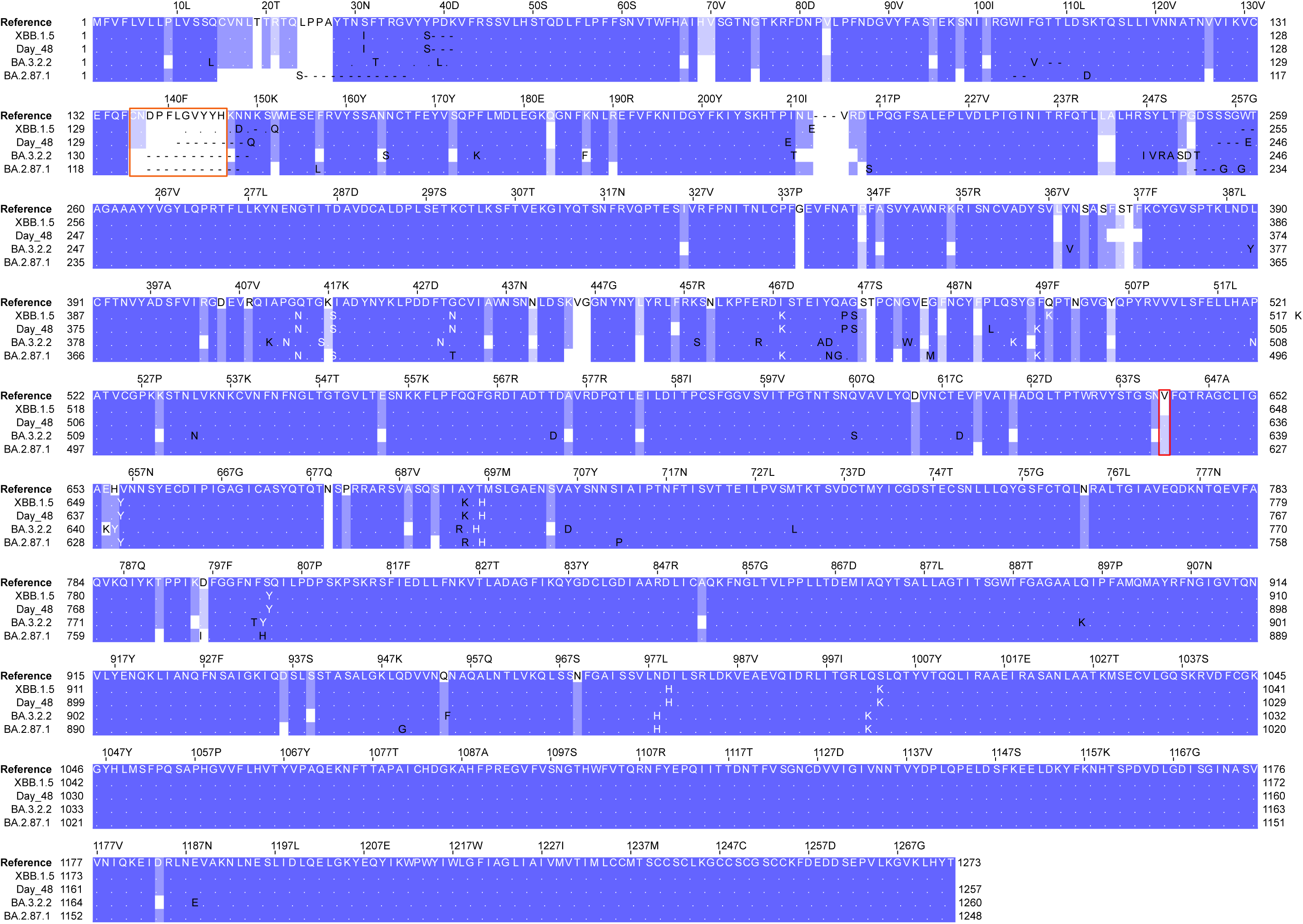
Multiple sequence alignment of Spike proteins from the within-host virus and divergent SARS-CoV-2 lineages. MUSCLE alignment of Spike sequences from Wuhan-Hu-1, XBB.1.5, the day 48 within-host sample, BA.3.2.2 (EPI_ISL_20282378), and BA.2.87.1 (EPI_ISL_18849991). The orange box marks the NTD region encompassing the 138-145 deletion. The red box marks residue 642, and V642G is present in the day 48 sequence, BA.3.2.2, and BA.2.87.1 but not in XBB.1.5. NTD, N-terminal domain.

**FIG S2.**
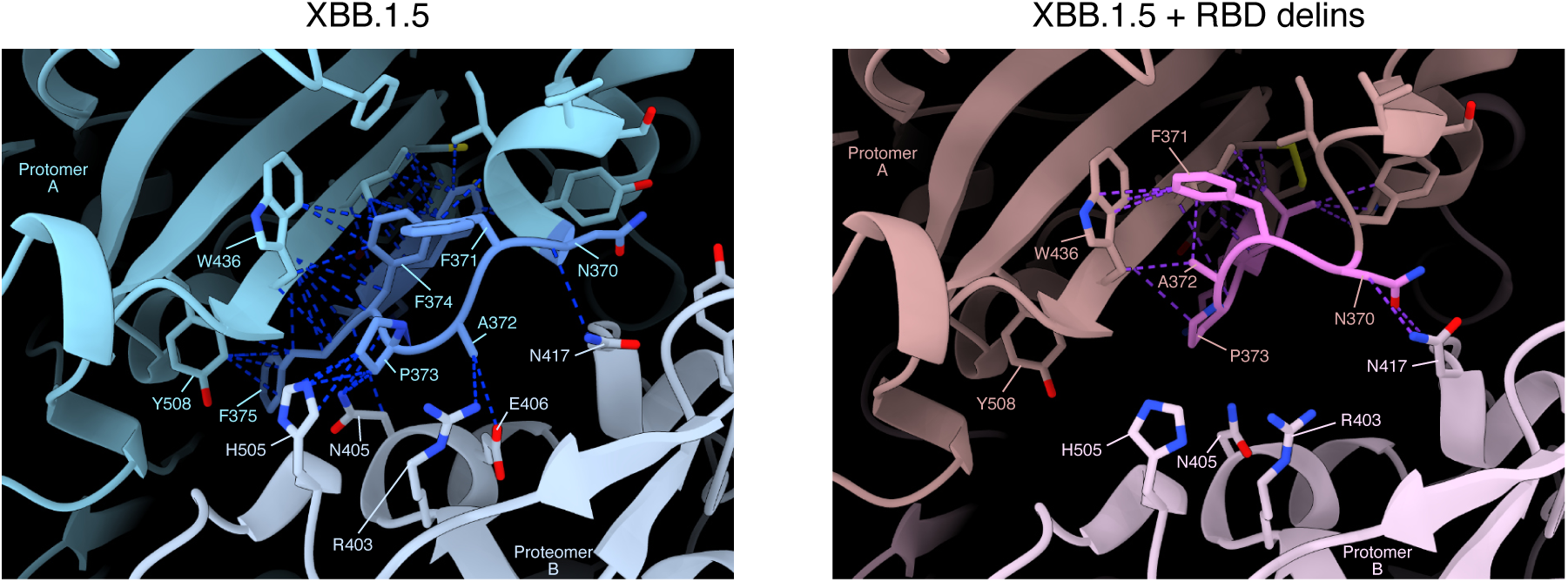
Predicted effects of the RBD delins on local contacts. Close-up views of AlphaFold 3-predicted Spike trimers for XBB.1.5 (A) and XBB.1.5 containing the RBD delins (B), centered on the altered RBD region near the interface between chain A and the adjacent chain B. Protein backbones are shown as cartoons and selected residues as sticks. Dashed lines represent local atomic contacts identified in UCSF ChimeraX from van der Waals overlap. The model containing the RBD delins shows an apparent reorganization and reduction of local contacts relative to XBB.1.5. Residue numbering follows the WA1 reference sequence. These models provide a qualitative structural hypothesis and do not establish the effect experimentally. RBD, receptor-binding domain.

**FIG S3.**
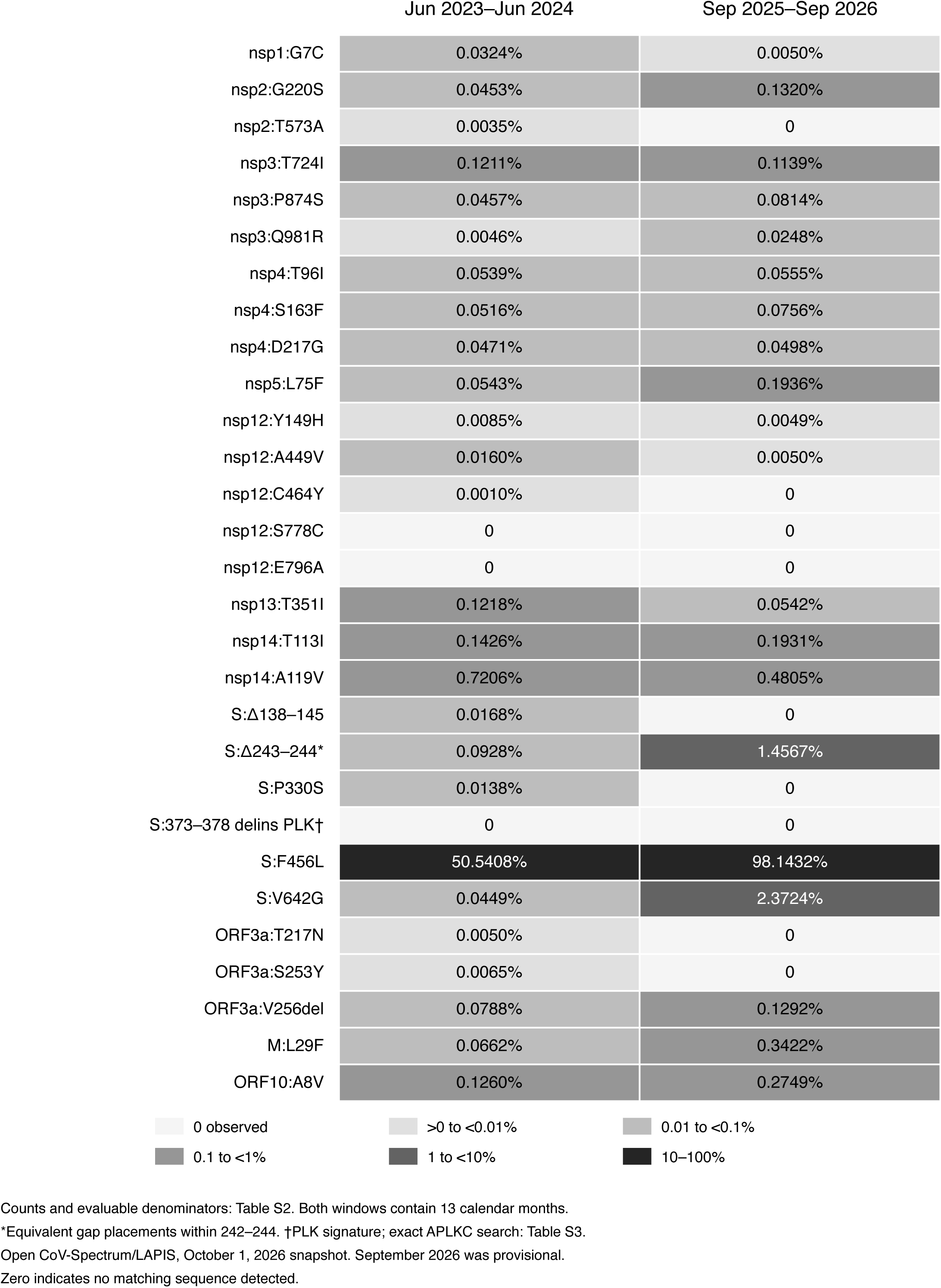
Public US sequence frequencies. Frequencies of the 29 changes shown in Figure 3 among evaluable US consensus sequences collected during June 1, 2023–June 30, 2024 and September 1, 2025–September 30, 2026. Labels show percentages and shading indicates frequency bands. Counts, denominators, and query definitions are provided in Table S2. The RBD PLK overview signature differs from the exact anchored APLKC search in Table S3. September 2026 data were provisional.

**FIG S4.**
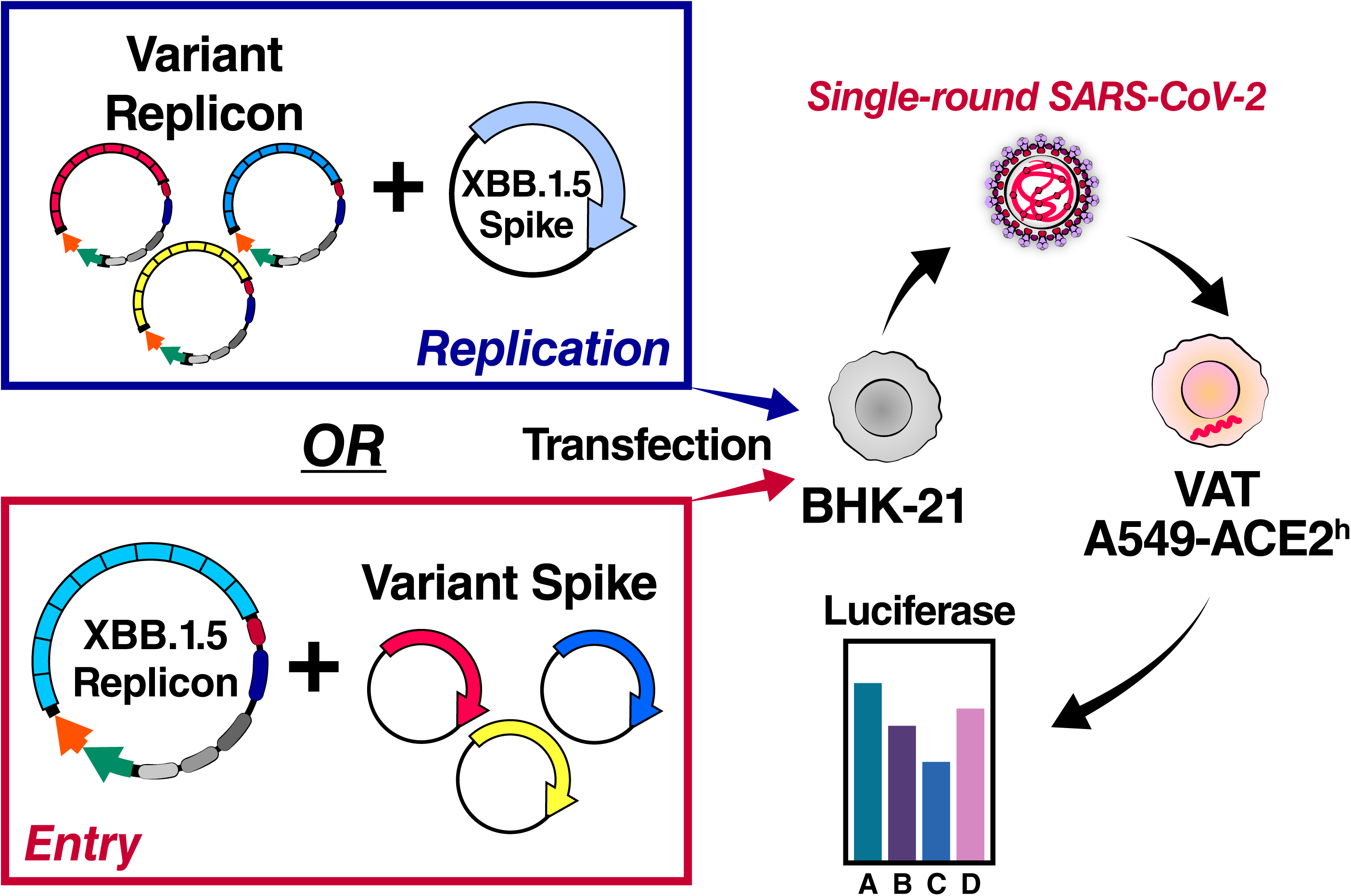
Design of the single-round SARS-CoV-2 replicon assays. Schematic provided by Taha Y. Taha showing how Spike and replicon variation were evaluated separately. For entry comparisons, the XBB.1.5 replicon was held constant while the Spike expression plasmid was varied. For RNA-replication comparisons, XBB.1.5 Spike was held constant while the replicon was varied. Particles produced in BHK-21 cells were used for single-round infection of VAT or A549-ACE2 recipient cells, with luciferase as the readout. Construct compositions are listed in Table S1.

## SUPPLEMENTAL TABLE LEGENDS

**TABLE S1. SARS-CoV-2 Spike and replicon construct compositions used in the functional assays.**

XBB.1.5 backbone mutations relative to Wuhan-Hu-1 and the patient-derived changes included in each Spike expression plasmid and Spike-deficient replicon are listed.

**TABLE S2. Frequencies of the changes featured in Figure 3**

Counts, evaluable denominators, and percentages for the 29 changes shown in Figure 3 in the two US collection periods. Numbering follows Wuhan-Hu-1, with gene coordinates mapped to the API annotations. Single-site queries used coverage at the relevant position, whereas linked patterns required calls across the queried interval and its flanks. Known deletion gaps were allowed. Evaluable refers to consensus calls, not raw-read depth. No additional host, lineage, or global quality filters were applied. Zero indicates no matching sequence detected in the available dataset.

The Δ138–145 signature required retained flanks and no intervening insertion. The Δ243–244 signature allowed equivalent placements of one leucine and two gaps within residues 242–244, with retained flanks and no insertion. The RBD overview signature required PLK in order across residues 373–378, allowing all 20 placements of three gaps, retained flanks, and no insertion, and its denominator required amino-acid calls across residues 372–379. ORF10 A8V was queried as a valine codon at nucleotides 29579–29581, with calls required at all three bases.

**TABLE S3. Exact and related RBD products**

Part A reports worldwide exact APLKC counts. This search required retained A372 and C379, allowing alternative gap placements within the intervening interval and checking separate insertion annotations. Evaluable records had amino-acid calls across residues 372–379 and nucleotide calls across bases 22676–22699, including known gaps.

Part B reports US exact APLKC, related APXKC (X denotes any standard amino acid), and constrained deletion/insertion searches. Candidate nucleotide sequences were checked for an intact local reading frame and agreement with the protein product. The constrained deletion screen required 1–4 deleted positions within residues 374–378, with residues 373 and 379 retained. Insertions of 1–4 amino acids, alone or with deletions, and exploratory five-amino-acid insertions were also screened. All-date rows include the dated windows, and categories may overlap.

Part C lists the four APFKC records, all collected in 2022: BA.1 (one), BA.1.1 (one), and BA.1.15 (two). Their local product contains phenylalanine at the position corresponding to leucine in the patient’s APLKC product. Their deletions extend outside residues 374–378, so they are excluded from the constrained deletion category. Hyphens denote alignment gaps.

